# Introducing PIGMO, a novel PIGmented MOuse model of Parkinson’s disease (V1)

**DOI:** 10.1101/2025.05.09.653000

**Authors:** J Chocarro, S Marana, M Espelosin, AJ Rico, G Ariznabarreta, E Lorenzo-Ramos, MM Ilarduya, R Hernandez-Alcoceba, M Chillón, M Vila, JH Kordower, AHV Schapira, A Garcia-Osta, M Cuadrado-Tejedor, JL Lanciego

**Affiliations:** CNS Gene Therapy Program, Center for Applied Medical Research (Cima), University of Navarra, 31008 Pamplona, Spain; Centro de Investigación Biomédica en Red de Enfermedades Neurodegenerativas (Ciberned-ISCIII), 28031 Madrid, Spain; Aligning Science Across Parkinson’s (ASAP) Collaborative Research Network, Chevy Chase, MD, USA; Institut des Neurociences, Department of Biochemistry and Molecular Biology, Universidad Autónoma de Barcelona; Barcelona, Spain; Val d’Hebron Research Institute, Barcelona, Spain; Institució Catalana de Recerca i Estudis Avançats (ICREA), Barcelona, Spain; ASU-Banner Neurodegenerative Disease Research Center, Arizona State University, Tempe, AZ, USA; Department of Clinical Neurosciences, University College London (UCL) Institute of Neurology, Royal Free Campus, Rowland Hill Street, London, UK

**Keywords:** neuromelanin, alpha-synuclein, Lewy bodies, substantia nigra, neurodegeneration

## Abstract

There is a pressing need for the development, characterization, and standardization of animal models of Parkinson’s disease (PD) that properly mimic the cardinal features of this disorder, comprising both the motor phenotype and neuropathological signatures. In the past few years, animal modeling has moved from neurotoxin-based approaches toward viral vectors carrying a given genetic payload of interest. Here, to induce pigmentation of the mouse brain upon systemic delivery, we took advantage of a modified adeno-associated viral vector capsid engineered to bypass the blood-brain barrier and coding for the human tyrosinase gene (AAV9-P31-hTyr). Obtained results revealed an ongoing pigmentation of catecholaminergic centers related to the pathophysiology of PD, such as the substantia nigra pars compacta, ventral tegmental area, and locus coeruleus. Moreover, pigmented dopaminergic neurons exhibited Lewy body-like intracytoplasmic inclusions, a progressive nigrostriatal degeneration, and a time-dependent PD motor phenotype. The bilateral pigmented mouse model of PD generated this way is highly reproducible, does not require stereotaxic surgery for viral vector deliveries, and opens unprecedented possibilities for preclinical testing of therapeutic candidates designed to reduce disease progression rates.

## INTRODUCTION

In the past few decades, the field of Parkinson’s disease (PD) has witnessed the introduction of novel generations of animal models that played an instrumental role in advancing the current understanding of the underlying physiopathological mechanisms and the development of therapeutic candidates. However, important limitations remain to be addressed, since by definition, any given model only replicates a small fraction of the inherent complexity of the clinicopathological spectrum of PD, and therefore, in most cases, models themselves do not accurately represent the real disease (Zeiss et al., 2017; Ransohoff, 2018; Marmion and Kordower, 2018; Baker and Björklund, 2020; Shadrina and Slominsky, 2021; Sturchio et al., 2024). Nevertheless, it is undeniable that available animal models of PD have paved the way for the implementation of pharmacological and neurosurgical therapies, providing patients with a clear benefit, both in terms of symptomatic alleviation and improvement of quality of life. Despite all the successes achieved with symptomatic treatments, disease-modifying therapeutics tested in existing models repeatedly failed when translated into clinical trials, and thus, rather than the putative candidate, it is very likely that the main limitations lie within the animal models themselves (Ransohoff, 2018; Chocarro and Lanciego, 2025).

Although rodent models based on viral vector-mediated expression of different alpha-synuclein species (wild type or mutated) have provided promising results by overcoming existing barriers in earlier generations of neurotoxin-based models and transgenic mice, these models also have their inherent limitations (Marmion and Kordower, 2018; Bezard et al., 2025). To mention a few, alpha-synuclein expression obtained this way is far beyond physiological levels and often results in a variable degree of cell loss and nigral pathology. Choices made regarding the viral serotype, promoter, and delivery route directly influence the obtained phenotype; therefore, a better standardization is still needed to ensure appropriate outcomes and reproducibility (Chocarro and Lanciego, 2025).

The introduction of adeno-associated viral vectors (AAVs) coding for the human tyrosinase gene (hTyr) as tools for modeling PD in rats and non-human primates (Carballo-Carbajal et al., 2019; Chocarro et al., 2023) has opened a new avenue in the field of animal modeling. The AAV-mediated enhanced expression of hTyr leads to a time-dependent accumulation of neuromelanin (NMel), a progressive death of pigmented dopaminergic neurons, the presence of Lewy bodies made of endogenous alpha-synuclein, and a pro-inflammatory environment, together with a motor phenotype (the latter being observed only in rats and not in macaques). Moreover, a transgenic mouse based on the constitutive expression of hTyr has been made available recently (Laguna et al., 2024). Animal models of PD obtained in these ways are based on earlier postulates linking NMel pigmentation and dopaminergic cell vulnerability (Hirsch et al., 1988, 1989; Kastner et al., 1992) as well as on accumulated clinical evidence showing a bi-directional relationship between the incidence of PD and melanoma, a cancer of skin melanocytes (reviewed in Flores-Torres et al., 2024). Indeed, alpha-synuclein has been detected in cultured melanoma cells and tissues derived from patients with melanoma (Dean and Lee, 2021). Clinical associations between Gaucher disease and melanoma have also been reported (Kramer et al., 2016; Murugesan et al., 2018). In this regard, mutations in the GBA1 gene (encoding the lysosomal enzyme glucocerebrosidase) currently represent the main genetic risk factor for PD (Blandini et al., 2019).

Since mice are the most commonly used experimental laboratory animals, we took advantage of an AAV serotype 9 capsid engineered to bypass the blood-brain barrier (BBB) upon intravenous administration (Nonnenmacher et al., 2020; Bunuales et al., 2024). BBB transcytosis of AAV9-P31 capsid variant is mediated by the carbonic anhydrase IV receptor, a protein expressed on the surface of endothelial cells (Shay et al., 2023). The intravenous administration of AAV9-P31 coding for the hTyr gene resulted in a novel rodent model (termed PIGMO model) mimicking the known motor and neuropathological phenotypes of human PD with unprecedented accuracy.

## MATERIALS AND METHODS

### Study design

This study aims to develop a novel pigmented mouse model of PD (PIGMO) that would enhance existing knowledge of underlying mechanisms leading to dopaminergic neurodegeneration, further increasing the chances of success when testing disease-modifying therapeutic candidates. The goal was to generate an accessible mouse model (e.g., fully reproducible, progressive, predictable, and without the inherent biases of stereotaxic surgeries) that mimics the neuropathological signatures of this disorder to the best possible extent, together with the characteristic progressive motor phenotype. Accordingly, we sought to evaluate whether the systemic delivery of a BBB-penetrant AAV9 capsid variant coding for the human tyrosinase gene (AAV9-P31-hTyr) can induce a bilateral, time-dependent pigmentation of catecholaminergic brain centers related to PD pathophysiology such as the substantia nigra pars compacta (SNpc), ventral tegmental area (VTA) and locus coeruleus (LC). In keeping with earlier data obtained upon intranigral deliveries of AAV1-hTyr (Carballo-Carbajal et al., 2019; Chocarro et al., 2023), Lewy body-like pathology and progressive cell death were expected to be found in catecholaminergic neurons of the CNS (in particular, within dopaminergic neurons of the SNpc). Adult mice were injected in the retro-ocular vein plexus with AAV9-P31-hTyr or an inactivated construct for control purposes (AAV9-P31-(i)hTyr). To delineate a timeline, injected mice were euthanized at 1, 4, 8, and 12 months post-viral deliveries, and motor behavior was assessed with rotarod and catalepsy tests before sacrifice. Brain tissue samples were processed for histological analysis, comprising intracellular neuromelanin levels, intracytoplasmic inclusions, and nigrostriatal damage.

### Experimental animals

Adult male and female mice (FVN/N strain; Charles River Laboratories, Barcelona, Spain) were used in this study. Animal handling and experimental procedures were conducted in accordance with the European Council Directive 2010/63/EU and in keeping with the Spanish legislation (RD 53/2013). The experimental design was approved by the Ethical Committee for Animal Experimentation of the University of Navarra (ref. CEEA 110-21) as well as by the Department of Animal Welfare of the Government of Navarra.

### Viral vector production

Recombinant AAV vector serotype 2/9 capsid variant P31 (Nonnenmacher et al., 2020) expressing the human tyrosinase cDNA driven by the CMV promoter (AAV9-P31-hTyr) and the corresponding control vector with an inactivated, scrambled human tyrosinase cDNA sequence [AAV9-P31-hTyr(i)] were produced at the Viral Vector Core Production Unit of the Autonomous University of Barcelona, Spain (UPV-UAB; https://www.viralvector.eu/). In brief, AAVs were produced by triple transfection of 2 x 108 HEK293 cells with 250 μg pAAV, 250 μg pRepCap, and 500 μg pXX6 plasmid mixed with polyethylenimine (Sigma-Aldrich). The UPV-UAB core facility generated a pAAV plasmid containing the inverted terminal repeats (ITRs) of the AAV2 genome, a multi-cloning site to facilitate cloning of expression cassettes, and an ampicillin resistance gene for selection. Two days after transfection, cells were harvested by centrifugation, resuspended in 30 ml of 20 mM NaCl, 2 mM MgCl_2_, and 50 mM Tris-HCl (pH 8.5), and lysed by three freeze-thawing cycles. Cell lysate was clarified by centrifugation, and the AAV particles were purified from the supernatant by iodixanol gradient as previously described (Zolotukhin et al., 1999). Next, the clarified lysate was treated with 50 U/ml of benzonase (Novagen; 1 h at 37 °C) and centrifuged. The vector-containing supernatant was collected and adjusted to 200 mM NaCl using a 5 M stock solution. To precipitate the virus from the clarified cell lysate, polyethylene glycol (Sigma-Aldrich) was added to a final concentration of 8%, and the mixture was incubated (3 h, 4 °C) and centrifuged. Pellets containing AAVs were resuspended in 20 mM NaCl, 2 mM MgCl_2_, and 50 mM Tris-HCl (pH 8.5) and incubated for 48 h at 4 C. The AAV titration method used was based on the quantitation of encapsulated DNA with the fluorescent dye PicoGreen®. Obtained vector concentrations were 1.16 x 10 gc/ml for AAV9-P31-hTyr and 1.76 x 10 gc/ml for AAV9-P31-hTyr(i). The plasmid map and sequence are provided in Supplementary Figure 1.

### Systemic deliveries of AAVs

Animals were anesthetized with isoflurane and treated with the analgesic buprenorphine (Brupex®; 0.1 mg/Kg). A drop of ophthalmic anesthetic (0.5% proparacaine hydrochloride) was also placed on the eye receiving the injection. Viral vectors were administered systemically through a retro-orbital injection of the venous sinus according to Yardeni et al. (2011), a more humane approach than alternative intravascular deliveries, and commonly used as the standard choice for viral vector deliveries in mice (Mitchell et al., 2000; Jerebtsova et al., 2005; Deverman et al., 2016; Knezevic et al., 2016; Gruntman et al., 2017; Prabhakar et al., 2021; Grodem et al., 2023; Okamoto et al., 2023; Tang et al., 2024; Li et al., 2025). Delivery was performed in the right retro-orbital sinus (right-handed operator) with a 30-gauge, 0.5-in insulin needle and syringe (Omnican®; ref. 9151117S; Braun, Melsungen, Germany), comprising a total volume of 100 μL made of 50 μL of AAVs diluted in 50 μL of PBS. The AAV suspension was slowly and smoothly injected. Once the injection was completed, the needle was slowly withdrawn to minimize reflux through the injection site. Experimental subjects were divided into four groups, each made of one control specimen and three treated animals.

### Necropsy and tissue processing

Animals were anesthetized with ketamine/xylazine (80/10 mg/Kg) and perfused transcardially with a saline solution (0.9% NaCl; 5 min; 9.5 ml/min), followed by 25 ml of a 4% paraformaldehyde-buffered solution (PFA). Once perfusion was completed, the brains were removed from the skull and post-fixed in 4% PFA for 24 h and then stored in a cryoprotectant solution containing 20% glycerin and 2% DMSO in neutral PBS. Next, frozen coronal sections (40 μm-thick) were obtained in a sliding microtome (Microm HM-400) and collected in 0.125 M PBS pH 7.4 as 10 series of adjacent sections. These series were used for (i) direct neuromelanin visualization; (ii) immunoperoxidase detection of TH; (iii) Masson-Fontana stain; (iv) cytoarchitectural stain with Neutral Red; (v) proteinase K-pretreated, triple immunofluorescent detection of alpha-synuclein (α-Syn), P62, and tyrosine hydroxylase (TH), combined with brightfield visualization of neuromelanin (NMel); (vi) proteinase K-pretreated, triple immunofluorescent detection of phosphorylated α-Syn(P-Ser 129), P62, and TH, combined with brightfield visualization of NMel; and (vii) proteinase K-pretreated, triple immunofluorescent detection of phosphorylated α-Syn(P-Ser 129), P62, and ubiquitin, combined with brightfield visualization of NMel. The remaining series of sections was stored at −80 C until further use, if required. A detailed protocol for necropsy and tissue processing is available elsewhere (Garcia-Gomara et al., 2025).

A complete list of the used primary and bridge antisera (biotinylated or Alexa® dye-conjugated) is provided in Table 1 below.

**Table 1.** List of antibodies and reagents.

| Antibody | Dilution | Incubation time | Source | Identifier | RRID |
| --- | --- | --- | --- | --- | --- |
| Goat anti-tyrosine hydroxylase (polyclonal) | 1:500 | Overnight | Abcam | Cat# ab101853 | AB_10710873 |
| Mouse anti-α-synuclein (monoclonal) | 1:40 | Overnight | Leica | Cat# NCL-L-ASYN | AB_442103 |
| Mouse anti-α-synuclein P-Ser129 (monoclonal) | 1:1000 | Overnight | Fujifilm Wako | Cat#015-25191 | AB_2537218 |
| Guinea Pig anti-P62 (polyclonal) | 1:1000 | Overnight | PROGEN | Cat# GP62-C | AB_2687531 |
| Mouse anti-ubiquitin (monoclonal) | 1:500 | Overnight | Abcam | Cat# ab7254 | AB_305802 |
| Donkey anti-Goat (Biotin-SP) | 1:600 | 120 min | Jackson | Cat# 705-065-147 | AB_2340397 |
| Donkey anti-Goat (Alexa488) | 1:200 | 120 min | Molecular Probes | Cat# A-11055 | AB_2534102 |
| Donkey anti-Goat (Alexa350) | 1:200 | 120 min | Molecular Probes | Cat# A-21081 | AB_141521 |
| Donkey anti-Goat (Alexa 633) | 1:200 | 120 min | Molecular Probes | Cat# A-21082 | AB_141493 |
| Donkey anti-Guinea Pig (Alexa594) | 1:200 | 120 min | Jackson | Cat# 706-585-148 | AB_2340474 |
| Donkey anti-Mouse (Alexa488) | 1:200 | 120 min | Molecular Probes | Cat# A-21202 | AB_141607 |
| Donkey anti-Mouse (Alexa546) | 1:200 | 120 min | Thermo Fisher | Cat# A-10036 | AB_11180613 |
| Donkey anti-Rabbit (Alexa488) | 1:200 | 120 min | Molecular Probes | Cat# A-21206 | AB_2535792 |

| Commercial assays | Dilution | Incubation time | Source | Identifier |
| --- | --- | --- | --- | --- |
| ABC Kit Standard |  | 60 min | Vector Labs | Cat# PK-4000 |
| Neutral Red | 0.40% | 1 min | Sigma | Cat# 72210 |
| Peroxidase chromogen V-VIP |  | 1 min | Vector Labs | Cat# SK-4600 |
| Proteinase K | 1 mg/ml | 10 min | Invitrogen | Cat# 25530049 |

### Analysis of pigmented neurons

Every 10th section was counterstained with Neutral Red and used to estimate the number of pigmented vs. non-pigmented neurons in the SNpc and VTA. Quantification was carried out through a dedicated, deep-learning bi-layered algorithm prepared with Aiforia® (www.aiforia.com; Penttinen et al., 2018) according to available protocol (https://dx.doi.org/10.17504/protocols.io.bp2l6xdwrlqe/v1). Five equally spaced coronal sections comprising the whole rostrocaudal extent of the SNpc and the VTA were sampled per animal. Sections were scanned at x20 in an Aperio CS2 scanner (Leica) and uploaded to the Aiforia cloud. The boundaries of the SNpc and VTA were outlined at low magnification (taking the exit of the 6_th_ cranial nerve as reference), and the algorithm was then used for quantifying the desired neuronal populations.

Furthermore, the location of three distinct neuronal profiles in the SNpc and in the VTA (TH+ / NMel+; TH+ / NMel-and TH-/NMel+) was carried out by camera lucida drawings made from three equally-spaced coronal sections at the level of the SNpc and VTA (comprising rostral, middle, and caudal midbrain levels) in each animal and for each time point.

### Quantification of intracellular neuromelanin levels

The intracellular density of NMel pigmentation was analyzed by measuring optical densitometry at the single-cell level with Fiji ImageJ software (NIH, USA). Obtained values were converted to a logarithmic scale according to the available protocol (Ruifrok and Johnston, 2001). The number of analyzed neurons within each region of interest (ROI) and at each time point is summarized in Table 2 below.

**Table 2.**
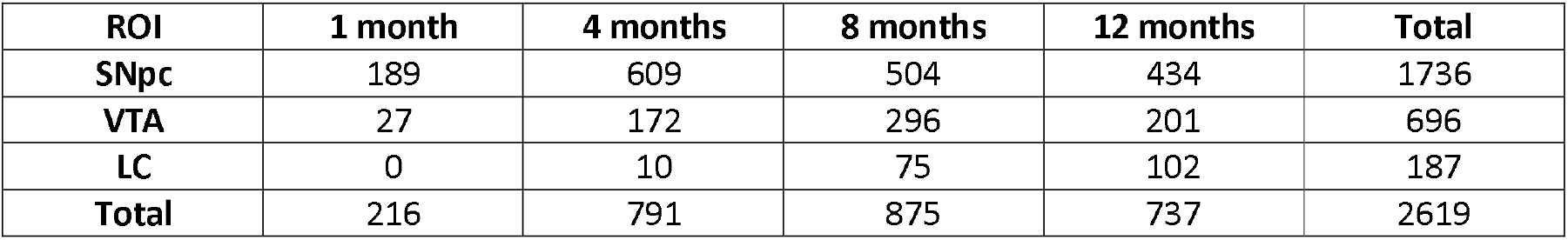
Number of neurons used for quantifying intracellular neuromelanin levels.

| ROI | 1 month | 4 months | 8 months | 12 months | Total |
| --- | --- | --- | --- | --- | --- |
| SNpc | 189 | 609 | 504 | 434 | 1736 |
| VTA | 27 | 172 | 296 | 201 | 696 |
| LC | 0 | 10 | 75 | 102 | 187 |
| Total | 216 | 791 | 875 | 737 | 2619 |

### Assessment of nigrostriatal damage

The second series of coronal sections was used for the immunohistochemical detection of TH (https://dx.doi.org/10.17504/protocols.io.8epv5x4wdg1b/v1). Sections rostral to the subthalamic nucleus (STN) were used for visualization of TH+ axon terminals at the level of the striatum using a standard diaminobenzidine peroxidase substrate (DAB), resulting in a dark brown precipitate. For each animal, TH optical intensities were measured in six equally spaced coronal sections of the striatum (pre- and post-commissural levels) with Fiji ImageJ software (NIH, USA), and the obtained values were converted to a logarithmic scale (Ruifrok and Johnston, 2001). Coronal sections caudal to the STN were used for the immunohistochemical detection of TH with purple peroxidase substrate (V-VIP; Lanciego et al., 1997, 1999). The V-VIP chromogen was chosen for TH detection in the SNpc, VTA, and LC since a better contrast is obtained for the NMel pigmentation against a purple background rather than the dark brown color that characterizes the DAB peroxidase substrate. The number of TH+ neurons in the SNpc and the VTA was quantified across six equally spaced coronal sections covering the whole rostrocaudal extent of the mesencephalon with a dedicated Aiforia® algorithm (https://dx.doi.org/10.17504/protocols.io.36wgq3x55lk5/v1).

### Neuronal inclusions

The presence of intracytoplasmic inclusions in pigmented catecholaminergic neurons of the SNpc, VTA, and LC was assessed under the confocal microscope by triple immunofluorescent detection of TH combined with characteristic markers of Lewy body pathology, such as P62 and phosphorylated alpha-synuclein (P-Ser129). One additional brightfield channel was included to visualize NMel pigmentation. Pre-digestion with proteinase K was conducted before immunofluorescent staining to remove soluble forms of alpha-synuclein. Moreover, another series of sections was used for triple immunofluorescent detection of phosphorylated alpha-synuclein, P62, and ubiquitin, to demonstrate that intracytoplasmic inclusions are ubiquitinated (Chocarro et al., 2023).

### Evaluation of motor phenotype

Standardized behavioral tests (rotarod and catalepsy tests; Garcia-Gomara et al., 2024, 2025) were used to evaluate motor readouts at different time-points post-administration of AAV9-P31-hTyr and AAV9-P31-hTyr(i) (one, four, eight, and twelve months). In brief, for the rotarod test, mice were placed on a revolving bar accelerating from 4 to 40 rpm over five minutes. To acclimate, animals were exposed for 30 seconds on day one and one minute on day two. On the test day, mice underwent two trials with a 30-minute interval, and results were expressed as the average latency to fall. The catalepsy test was used to measure muscular rigidity by placing the forepaws of the experimental subjects on a 4 cm-high bar. The time taken to adjust the posture was recorded across three trials, with a one-minute resting interval.

### Statistical analyses

Statistical analyses were performed in GraphPad Prism version 9.0.2 for Windows and Stata 14 (Stata Corp. 2017, Stata Statistical Software Release 15, College Station, TX; StataCorp LLC). Relevant tests are listed in the figure legends. Species with p <0.05 were considered statistically significant.

## RESULTS

### Time-dependent accumulation of neuromelanin

The systemic delivery of AAV9-P31-hTyr resulted in a time-dependent bilateral pigmentation of brain catecholaminergic centers of the mouse brain mimicking pigmentation patterns observed in NMel transgenic mice (Laguna et al., 2024). The conducted study was focused on the substantia nigra pars compacta (SNpc), the ventral tegmental area (VTA), and the locus coeruleus (LC), all these nuclei related to the pathophysiology of PD, according to the Braak hypothesis that postulates a clinico-pathological correlate of disease progression (Braak et al., 2002, 2003).

Initial traces of intracellular neuromelanin accumulation in SNpc neurons were observed as early as one month post-delivery of AAV9-P31-hTyr, with a moderate number of weakly pigmented neurons. By contrast, the same follow-up period resulted in a small number of pigmented neurons in the VTA, with minimal levels of intracellular neuromelanin. Accumulation of neuromelanin gradually increased over time, reaching levels high enough to enable a direct macroscopic visualization of the pigmented SNpc and VTA nuclei beyond four months post-viral vector deliveries (Figure 1). In parallel to the increase in neuromelanized SNpc neurons, intracellular pigmentation levels also rise over time. In other words, the number and NMel levels of pigmented neurons followed a time-dependent pattern (Figure 2). That said, although intracellular pigmentation levels within SNpc neurons still increased from eight to twelve months post-viral delivery, the total number of neuromelanized neurons showed some decline, likely reflecting a more pronounced nigrostriatal degeneration at twelve months. Moreover, a tier-specific pigmentation of dopaminergic neurons was observed upon the systemic delivery of AAV9-P31-hTyr in mice. Neurons in the ventral tier of the SNpc that are positive for aldehyde dehydrogenase type 1a1 (Aldh1a1+) are those accumulating NMel, whereby calbindin-positive neurons (CB+) in the dorsal tier never became pigmented (Supplementary Figure 2).

**FIGURE 1.**
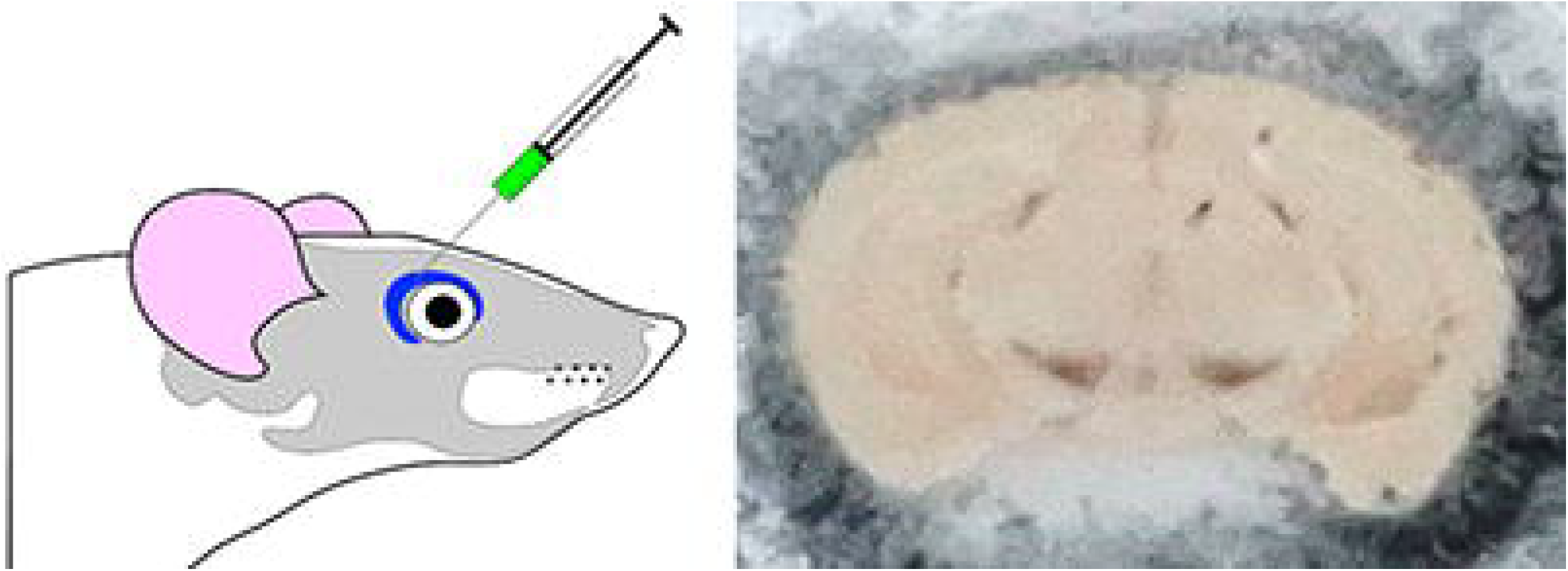
Systemic delivery of viral vectors. Schematic representation of the procedure for viral vector injections into the retro-ocular venous sinus of mice, together with a macroscopic visualization of the obtained neuromelanin pigmentation in the substantia nigra pars compacta (SNpc) and ventral tegmental area (VTA) as observed in a mouse injected with AAV9-P31-hTyr four months post-viral delivery.

**FIGURE 2.**
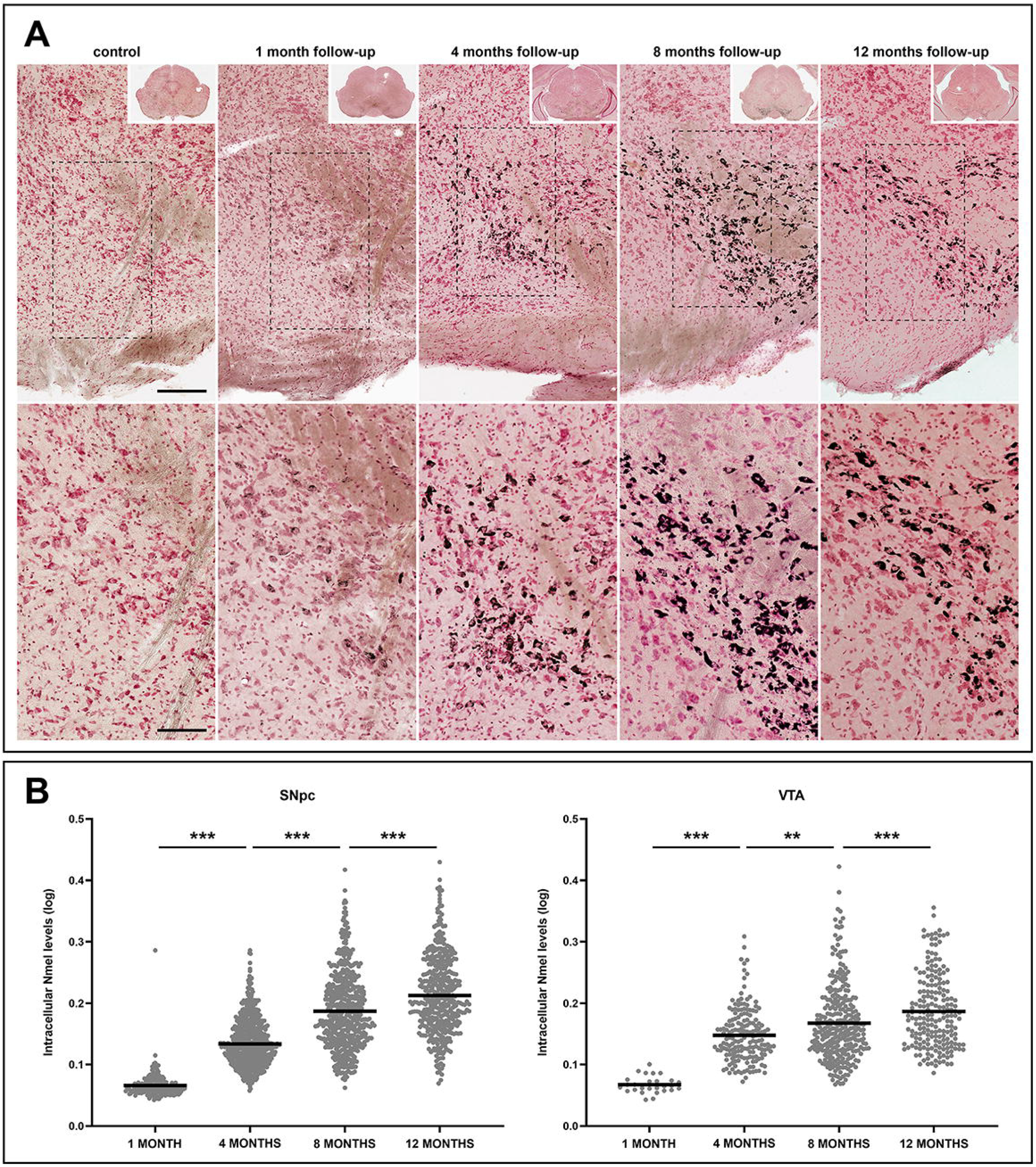
Time-dependent pigmentation of SNpc and VTA neurons. **(A)** Progressive accumulation of neuromelanin (NMel). A weak pigmentation was first observed one month post-injection of AAV9-P31-hTyr. Accumulation of NMel increased over time, as shown in representative photomicrographs taken four, eight, and twelve months of follow-up. Scale bars, 100 μm for low-magnification images and 200 μm in insets. **(B)** Box plots showing intracellular neuromelanin levels at the single-cell level for all experimental groups in the SNpc and VTA. In both structures, a significant augmentation of intracellular pigmentation was observed at all time points. Values are mean +/-SEM, nested ANOVA test with time as a fixed factor, and mice nested with fixed factor. At the SNpc level, p < 0.001 (four vs. one month, eight vs. four months, and twelve vs. eight months). p < 0.01 when comparing different four vs. eight months at the level of the VTA.

Compared to the SNpc, neurons in the VTA showed a similar temporal pattern of gradual pigmentation, but to a lower magnitude. The number of neuromelanized VTA neurons and their intracellular pigmentation levels are time dependent, however, the slope of said increase is far less pronounced than in the SNpc (Figure 2). Regarding the LC, noradrenergic neurons appeared to be more resistant to pigmentation. Neuromelanized neurons in the LC were first detected four months post-systemic delivery of AAV9-P31-hTyr, and exhibited minimal traces of intracellular NMel accumulation. Although more neurons in the LC became pigmented over time, no differences were observed when comparing intracellular pigmentation levels between four and eight months post-delivery. The highest accumulation of neuromelanin in the LC was found with a follow-up period of twelve months (Figure 3), reaching pigmentation levels only comparable to those observed in the SNpc and VTA with a survival time of four months (Figures 2 and 3).

**FIGURE 3.**
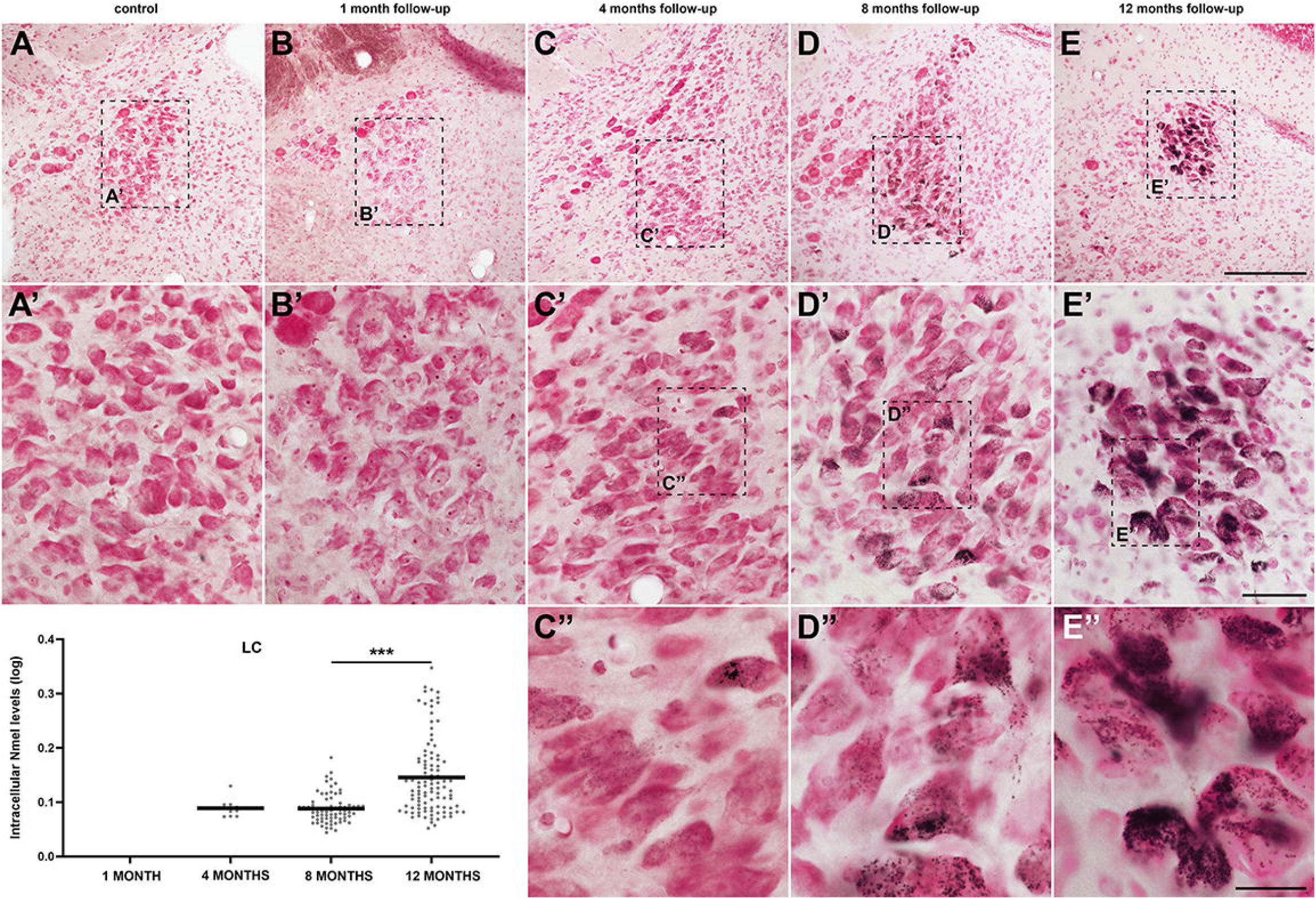
Time-dependent pigmentation of LC neurons. Noradrenergic neurons of the LC became progressively pigmented following the delivery of AAV9-P31-hTyr viral vector, although to a lower level than dopaminergic neurons in the SNpc and VTA. By twelve months of follow-up, accumulation of NMel is similar to SNpc and VTA neurons four months post-viral injection. Box plots indicate values for intracellular NMel pigmentation, represented as mean +/-SEM, nested ANOVA test with time as a fixed factor, and mice nested with fixed factor. p < 0.001 between eight and twelve months. Scale bars, 200 μm in panels A-E; 50 μm in A’-E’, and 20 μm in C’’-D’’.

### Lewy body pathology in pigmented neurons

The presence of intracytoplasmic inclusions was analyzed at different time points post-systemic delivery of AAV9-P31-hTyr. In keeping with earlier data linking NMel levels beyond a given threshold as drivers inducing Lewy body-like pathology (LBs; Carballo-Carbajal et al., 2019; Chocarro et al., 2023; Laguna et al., 2024), intracytoplasmic inclusions were observed in the SNpc of all animals with a follow-up period beyond 4 months (Figure 4). Although pigmented neurons were first noticed one month post-injection of AAV9-P31-hTyr, NMel levels were not high enough to trigger the accumulation of endogenous alpha-synuclein in the form of LBs after such a short survival time. LBs were first observed in dopaminergic neurons of the SNpc four months post-delivery of the BBB-penetrant viral vector encoding the human tyrosinase gene. Although the total number of LBs was not quantified, intracellular inclusions were more abundant in all animals sacrificed after eight and twelve months post-viral deliveries. Observed inclusions were positive for traditional markers of LB pathology, such as PK-resistant phosphorylated alpha-synuclein (Ser129), P62, and ubiquitin (Figure 5). Regarding the VTA, LB-like pathology was most often observed after eight and twelve months of follow-up. Finally, intracytoplasmic inclusions were only found in the LC twelve months post-viral delivery (Supplementary Figure 3), likely reflecting lower pigmentation levels that typically characterize LC neurons, which are equivalent to the pigmentation levels found in the SNpc and VTA after a follow-up of four months (Figures 2 and 3).

**FIGURE 4.**
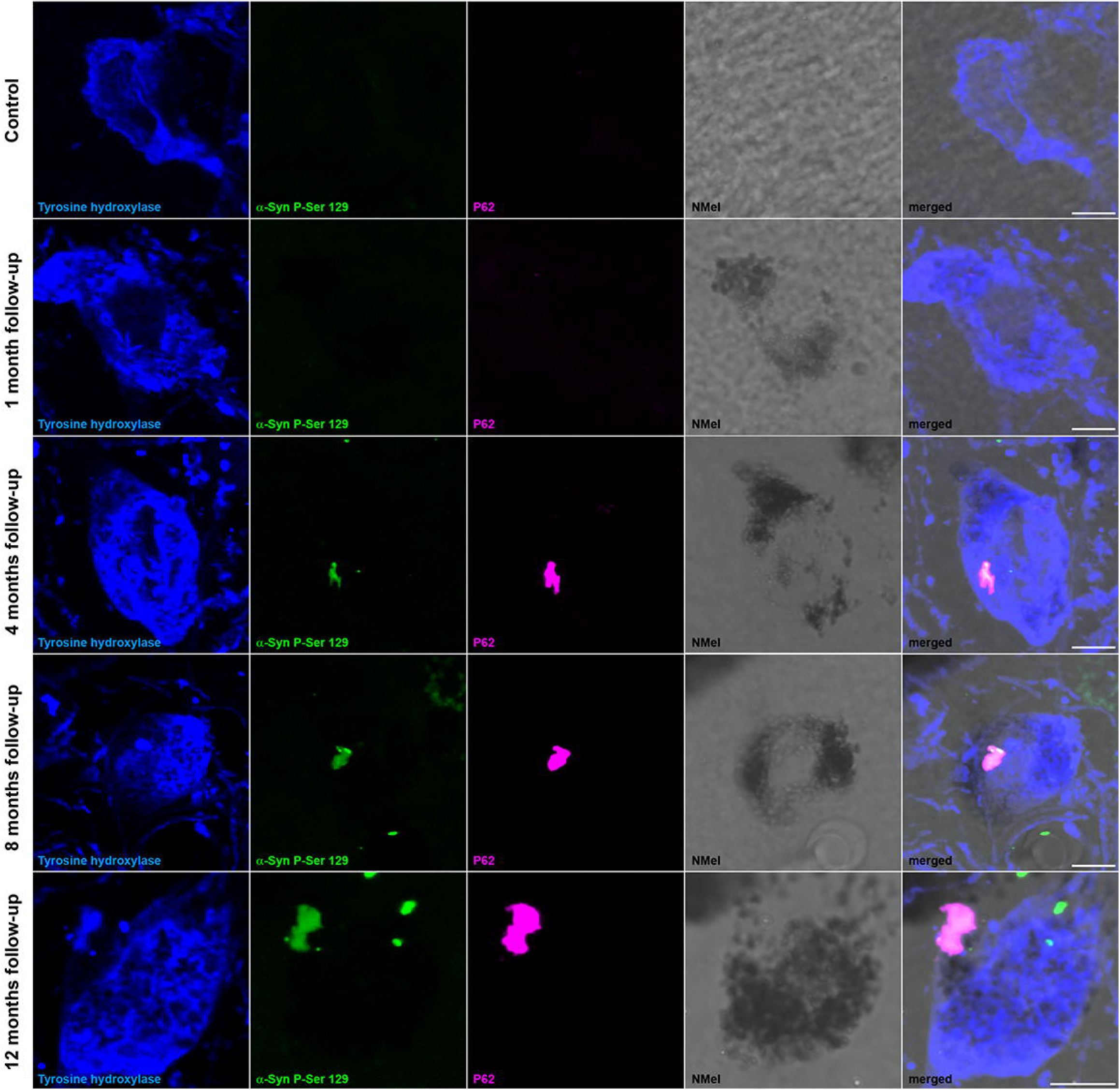
Intracytoplasmic aggregates in dopaminergic neurons. Intracellular inclusions within dopaminergic neurons of the SNpc (TH+; blue channel) followed a time-dependent pattern. Although a weak NMel accumulation is first found one month post-delivery of AAV9-P31-hTyr, pigmentation levels are not high enough to induce Lewy body-like intracytoplasmic aggregates. Inclusions positive for phosphorylated alpha-synuclein (P-Ser 129; green channel) and P62 (purple channel) are observed in experimental groups with four, eight, and twelve months of survival time, in parallel to the ongoing increase of pigmentation. Scale bars, 5 μm.

**FIGURE 5.**
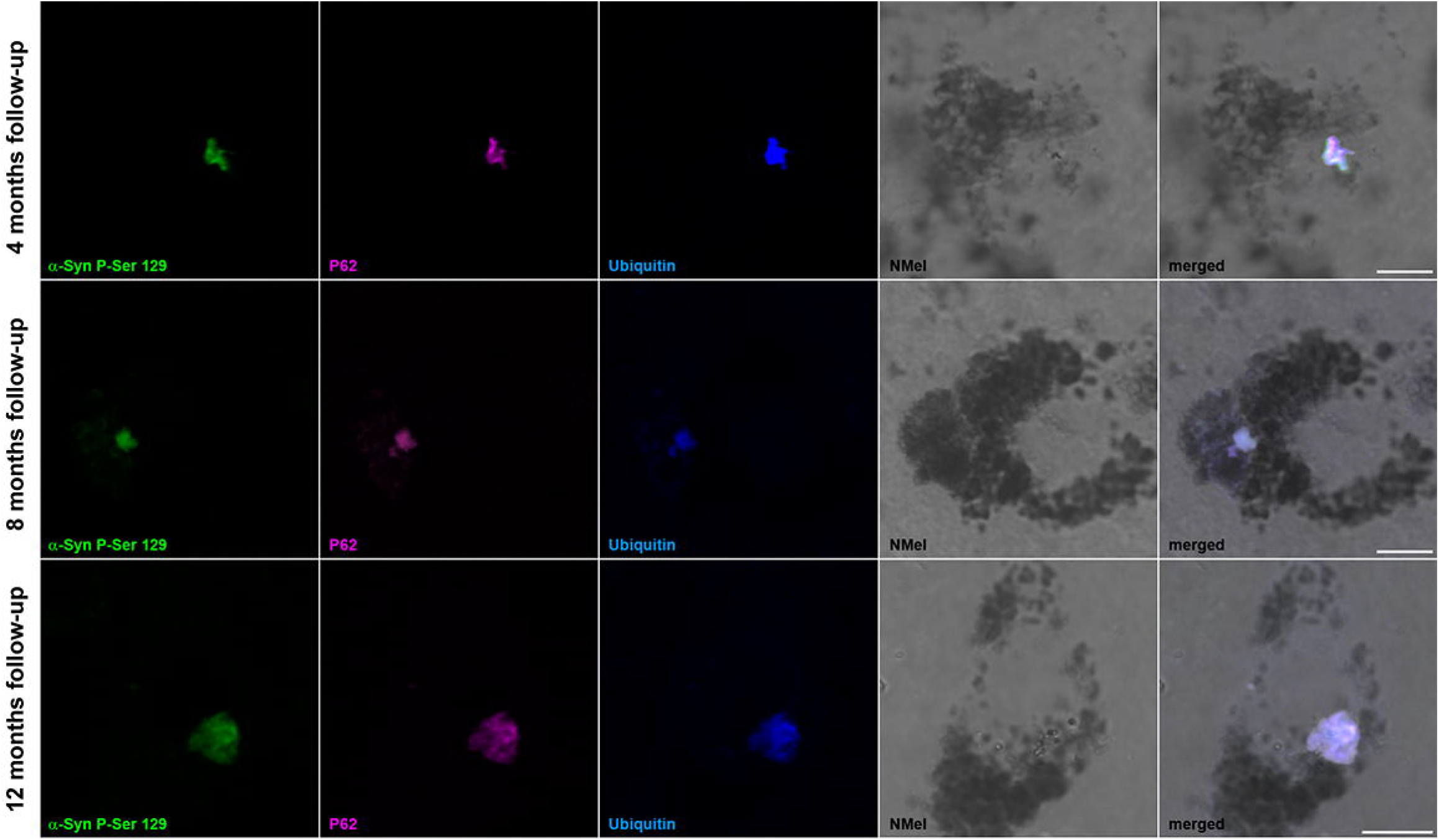
Markers of Lewy body-like inclusions in pigmented neurons of the SNpc. Triple immunofluorescent detection of Lewy body markers within pigmented neurons in the SNpc, comprising phosphorylated alpha-synuclein (P-Ser 129; green channel), P62 (purple), and ubiquitin (blue). As expected, intracytoplasmic inclusions were found in heavily neuromelanized neurons, bearing in mind that pigmentation beyond a given threshold is required to induce aggregation of endogenous alpha-synuclein in the form of Lewy bodies. Scale bars, 5 μm.

### Progressive nigrostriatal degeneration

In parallel to NMel accumulation and LB-like burden, the observed dopaminergic cell loss in the SNpc and the corresponding decline of dopaminergic terminals in the striatum followed a time-dependent pattern.

Optical densities for TH+ axon terminals in the striatum gradually declined over time (Figures 6A & B). Compared to control animals, no differences were observed one month post-delivery of the viral vector. A minor decrease was first observed at four months (10.34% of reduction in optical density) without reaching statistical significance. In animals euthanized eight months post-injection, a significant bilateral reduction in TH+ terminals was observed in dorsolateral territories of the striatum (34.48% of decline compared to control animals), further reaching a more severe reduction (50.12% on average) in animals with a follow-up of 12 months, where the loss of TH+ terminals in the dorsolateral striatum is far more pronounced.

**FIGURE 6.**
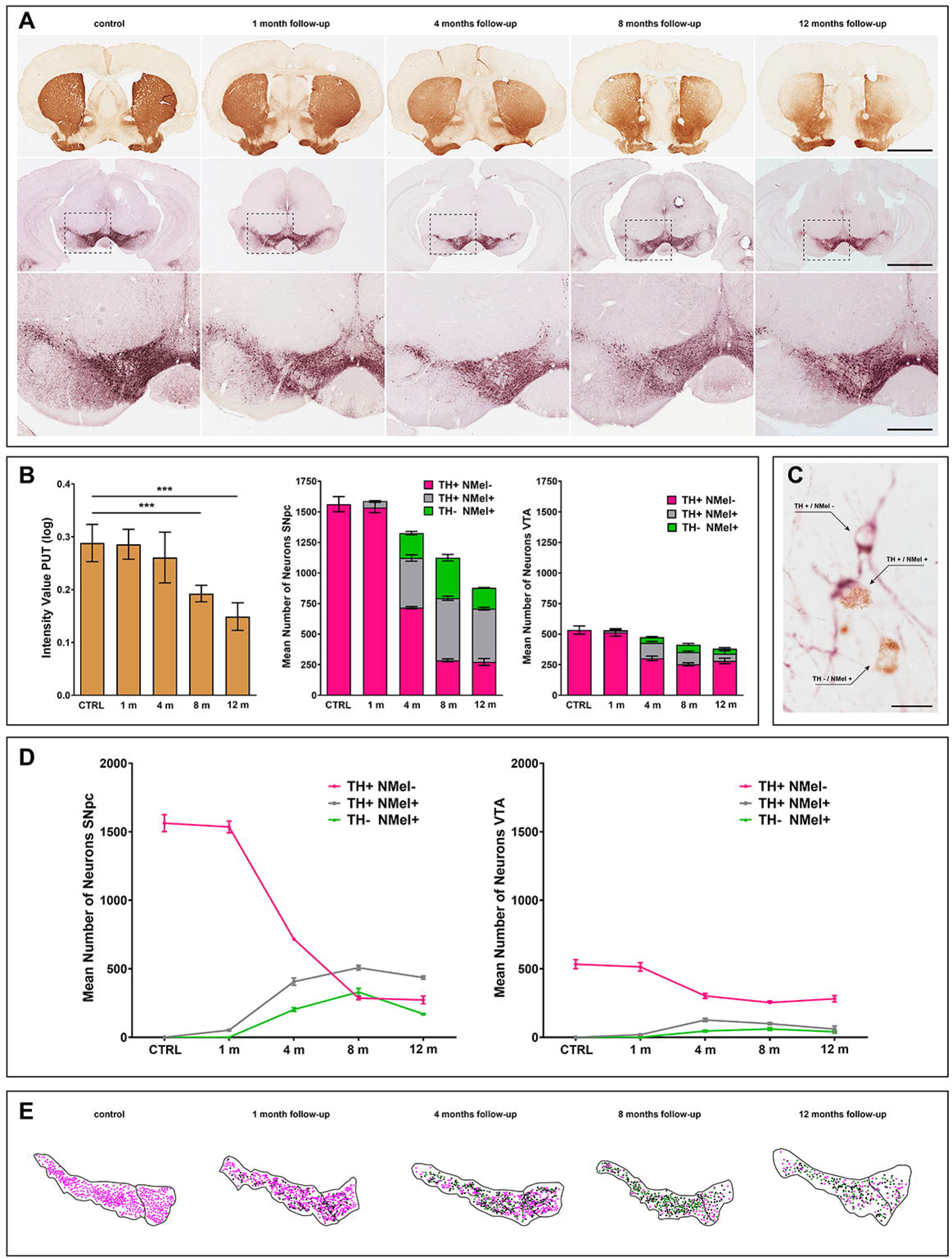
Progressive nigrostriatal lesion. The time-dependent NMel accumulation and the subsequent endogenous synucleinopathy triggered an ongoing lesion of the nigrostriatal pathway, both at origin (SNpc) and at destination (striatum). **(A)** Immunohistochemical detection of TH in the striatum (brown reaction product) and in the substantia nigra (purple peroxidase substrate) showing that nigrostriatal damage increased over time. A bilateral loss of TH+ terminals in dorsolateral striatal territories, reflecting progressive death of nigrostriatal-projecting neurons of the SNpc, was noticed, even at a low magnification. Scale bars, 2000 μm low-magnification images taken from the striatum and SNpc, and 500 μm in insets. **(B)** Histograms showing the progressive decline of TH+ terminals in the striatum and the corresponding phenotype-specific cell loss in the SNpc and VTA. The quantification of optical density for dopaminergic terminals in the striatum is represented as mean +/-SEM, unpaired t-test. p < 0.001 between control animals and eight months, and p < 0.0001 between controls and twelve months. **(C)** Photomicrograph taken from the SNpc in an experimental subject with eight months of follow-up showing three distinctive neuronal phenotypes, namely non-pigmented TH+ neurons (TH+ / NMel-), neuromelanized TH+ neurons (TH+ / NMel +), and ghost cells (TH-/ NMel+). Scale bar, 20 μm. **(D)** Schematic representation of the dynamic changes observed over time across the three characteristic cellular phenotypes. In the SNpc, the main cell loss was observed between one and four months, and from four to eight months. Beyond eight months of survival time, the number of TH+ / NMel-reached a plateau, likely suggesting the presence of a small cellular reservoir made of non-pigmented neurons that are more resilient to degeneration. Moreover, pigmented dopaminergic neurons (TH+ / NMel+) and ghost cells (TH-/ NMel+) followed a similar pattern, reaching a maximum peak at eight months and with a moderate decline at twelve months. This decline is higher for ghost neurons, which are the most vulnerable ones. **(E)** Camera lucida drawings taken from a representative level of the SNpc and VTA across all experimental groups showing the ongoing changes for all the different cellular phenotypes.

The gradual damage of TH+ terminals in the striatum was in keeping with a similar pattern of progressive dopaminergic cell loss observed in the SNpc and, to a minor extent, in the VTA (Figure 6). Compared to control specimens, the number of dopaminergic cells started to decline four months post-systemic deliveries of AAV9-P31-hTyr (16.67% of cell loss at four months), and this pattern of cell loss increased over time (27.88% at eight months, and 44.55% at twelve months). Of particular importance, dopaminergic cell death was found to be phenotype-dependent. Up to three distinct dopaminergic cell phenotypes were found, comprising (i) non-pigmented dopaminergic cells (TH+/NMel-), (ii) neuromelanized neurons (TH+/NMel+) and (iii) pigmented neurons that lost the TH phenotype (TH-/NMel+), the latter also known as “ghost cells” that are likely those that are more vulnerable to degeneration (Figure 6C). Pigmented dopaminergic neurons (TH+/NMel+) were observed as early as one month post-viral administration (Figures 2 & 6), although they represent a minimal fraction of the total number of dopaminergic neurons (3.12%). The number of TH+/NMel+ neurons increased at four months, when ghost cells were first observed. These two populations of pigmented neurons (TH+/NMel+ and TH-/NMel+) collectively accounted for 45.93% of the total number of dopaminergic neurons, meaning that by four months, roughly half of the SNpc neurons are pigmented. The ratio between pigmented and non-pigmented neurons increased at eight and twelve months (74.57% at eight months and 68.95% at twelve months). The most relevant decline in the number of TH+ neurons was observed from one to four months and from four to eight months. Beyond eight months, the number of TH+/NMel-neurons reached a plateau, likely meaning that non-pigmented neurons are more resilient to degeneration. The temporal dynamics for TH+/NMel+ and TH-/NMel+ neuronal phenotypes are roughly similar. These two cellular phenotypes reached a maximum peak at eight months, followed by a slight decline at twelve months (Figure 6).

Although all three different cellular phenotypes were also observed in the VTA, these neurons are far less vulnerable than those of the SNpc. Compared to the numbers of non-pigmented TH+ neurons, neuromelanized neurons (NMel+/TH+ & NMel+/TH-) barely represented half of the total dopaminergic cells in the VTA. Moreover, although a gradual cell loss was observed over time, such cellular neurodegeneration was smaller than in the SNpc (Figure 6).

### Motor phenotype

A time-dependent motor phenotype was observed in parallel to the extent of the nigrostriatal lesion. Compared to control animals, the experimental group comprising mice with a follow-up of 12 months exhibited a statistically significant difference in the rotarod test (Figure 7). Moreover, the catalepsy test resulted in a higher sensitivity, where significant differences were found in all experimental groups beyond four months post-delivery of AAV9-P31-hTyr (Figure 7). In summary and although each experimental group was made of a relatively low number of subjects, a clear time-dependent motor phenotype was observed with minimal inter-group differences.

**FIGURE 7.**
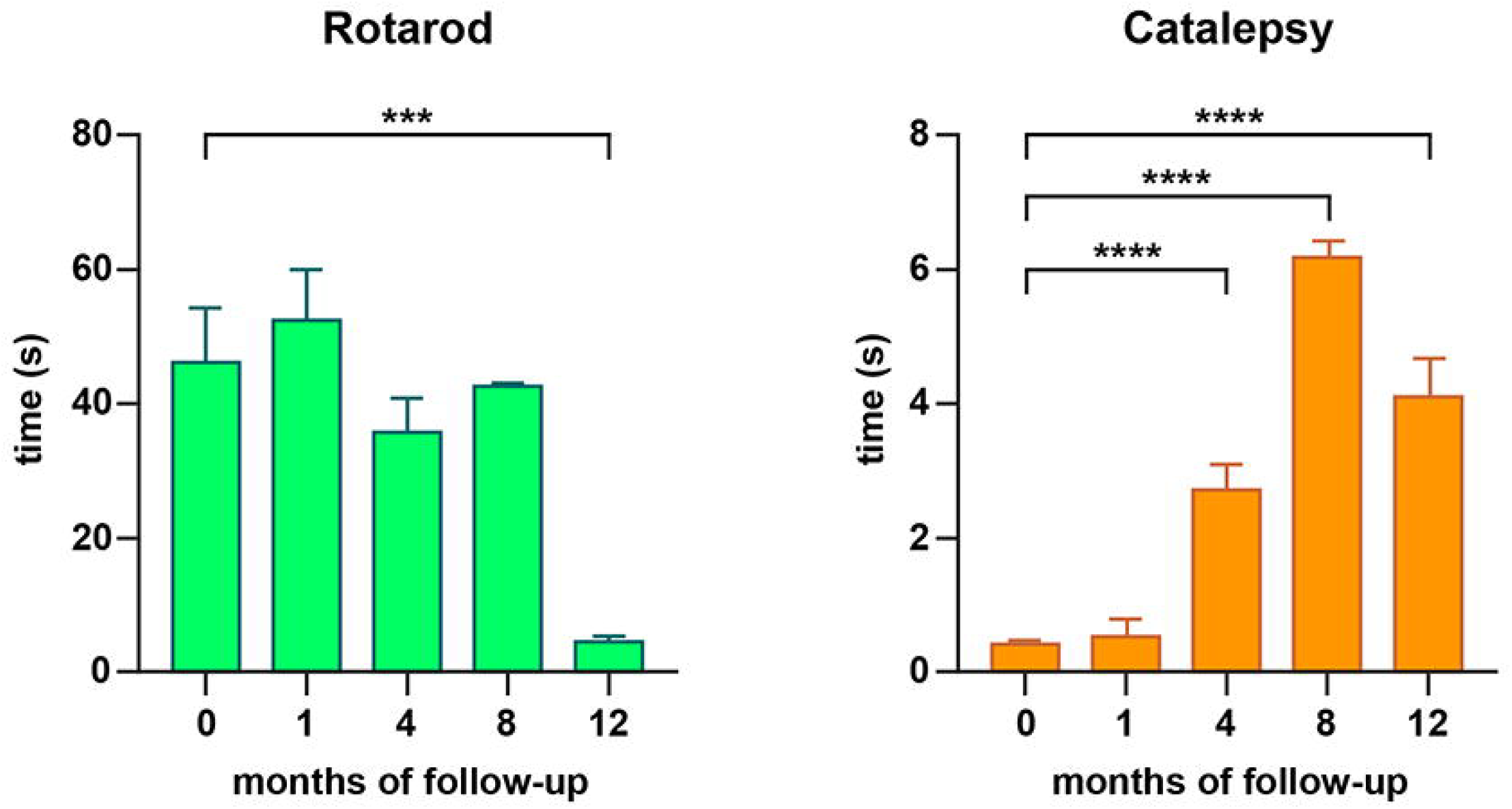
Motor readouts. The underlying nigrostriatal damage induced the appearance of a parkinsonian-like motor impairment as observed with the conducted motor evaluation. Statistically significant differences were found with the rotarod test at twelve months of follow-up compared to controls (p < 0.0001), whereby the catalepsy test was more sensitive in addressing a motor readout. Data are represented as mean +/-SEM, one-way ANOVA and Dunnett’s multiple comparison test, p < 0.001 was obtained for comparisons between control specimens and animals with four, eight, and twelve months of survival times.

### Peripheral organs

Since AAV9-P31-hTyr was delivered intravenously, samples from peripheral organs were taken at each time point to further address the potential presence of pigmentation. Analyses comprised paraffin-embedded tissue samples obtained from the lungs, heart, liver, spleen, small and large intestines, adrenal gland, kidney, striatal skeletal muscle, and gonads. Initial screenings were based on sections stained with H&E and non-stained sections. From all peripheral organs analyzed, high pigmentation levels were found in the red pulp of the spleen, beginning at four months post-viral delivery and gradually increasing over time (Supplementary Figures 4A-B). Furthermore, a very weak NMel accumulation was found in the sarcoplasmic cones of the cardiomyocytes at twelve months post-injection of AAV9-P31.hTyr (Supplementary Figures 4C-D). Pigmentation was never observed in peripheral organs other than the spleen and heart, irrespective of the follow-up time.

## DISCUSSION

Here, a novel mouse model of PD has been developed and characterized, mimicking the known motor and pathological signatures typically observed in human PD. The intravenous administration of AAV9-P31-hTyr induced (i) a bilateral, ongoing pigmentation of catecholaminergic neurons in the SNpc, VTA, and LC, (ii) a time-dependent Lewy body-like pathology in neuromelanized neurons, (iii) a progressive nigrostriatal degeneration, and (iv) a PD-related motor phenotype (Figure 8). Moreover, and in keeping with earlier animal models of enhanced neuromelanin pigmentation (Carballo-Carbajal et al., 2019; Chocarro et al., 2023; Laguna et al., 2024), the obtained data supported that neuromelanin accumulation beyond a given threshold triggers an endogenous synucleinopathy in the form of Lewy bodies.

**FIGURE 8.**
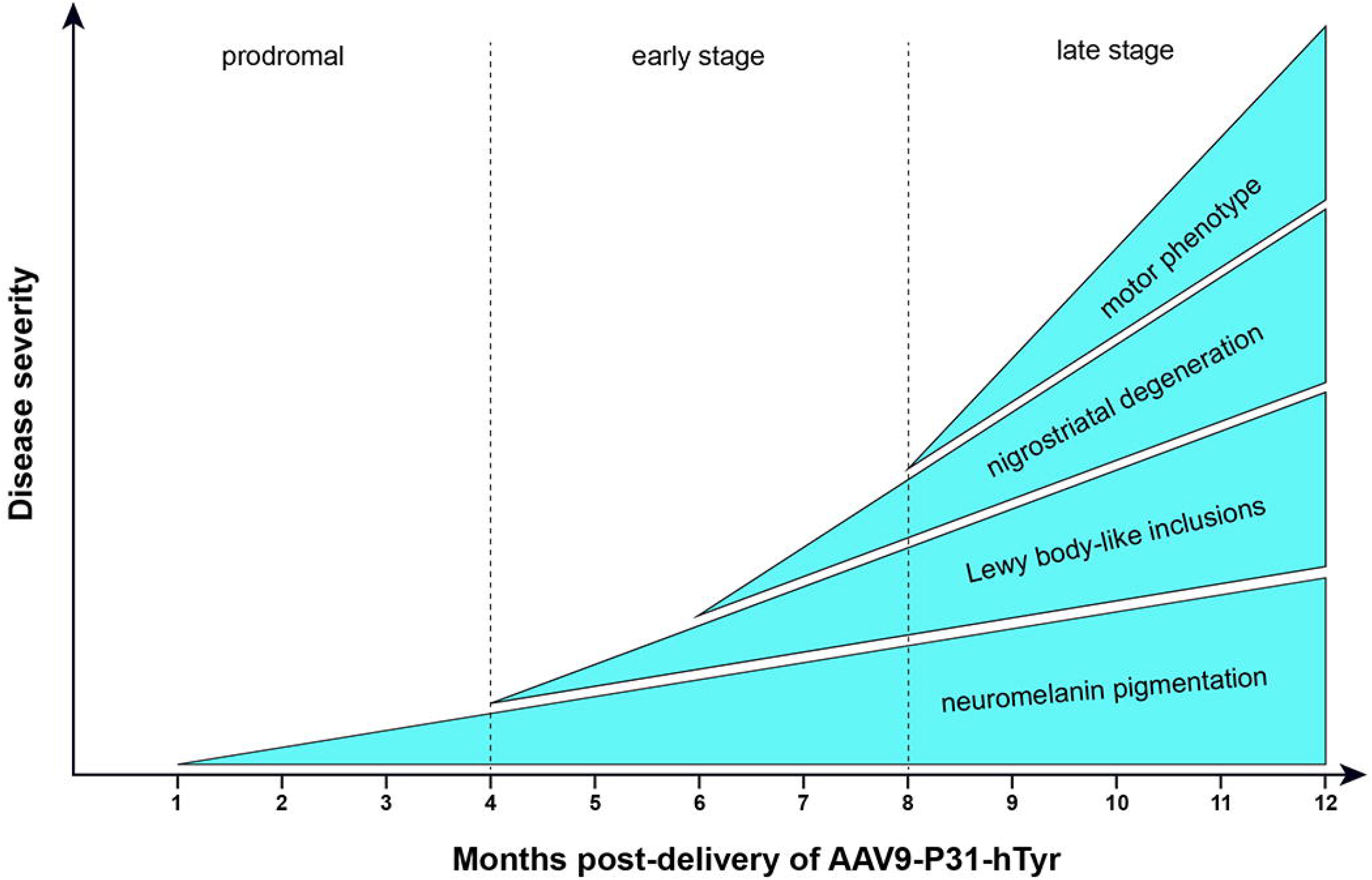
Outcomes of the PIGMO model. Schematic representation of the newly introduced PIGMO mouse model of Parkinson’s disease. The progressive cascade of events characterizing the PIGMO model defines well-characterized prodromal, early, and late stages, which would be instrumental for implementing novel disease-modifying therapeutic approaches. Since the model is predictable and has a wide therapeutic window, therapeutics can easily be administered once the underlying pathology has already started, and before reaching a non-returning point.

### BBB-penetrant viral capsid variants

The conducted study took advantage of a novel AAV9 capsid library designed by Voyager Therapeutics (Lexington, MA, USA; Nonnenmacher et al., 2020) using the so-called TRACER platform (tropism redirection of AAV by cell-type-specific expression of mRNA). AAV9 capsid variants were engineered to bypass the BBB by targeting the carbonic anhydrase IV receptor, a protein expressed on the surface of endothelial cells (Shay et al., 2023). Among the obtained capsid library, the P31 variant was chosen since it seems to be the best-performing one in terms of whole-brain transduction (Nonnenmacher et al., 2020). Upon intravenous administration, AAV9-P31 crossed the BBB and efficiently spread throughout the entire CNS, with a limited liver tropism. Neurons and astrocytes were transduced in the brain and spinal cord according to region-specific patterns (Bunuales et al., 2004).

There is a continuous race for the development of AAV capsid variants with the ability to bypass the BBB, in an attempt to transduce the brain and spinal cord. Although AAV9 has some degree of BBB penetrance, the first capsid specifically designed for this purpose was AAV2-PHP.B, and a subsequent variant known as AAV2-PHP.eB (Deverman et al., 2016). These capsids bypass the BBB by targeting the LY6A cellular receptor (Huang et al., 2019; Hordeaux et al., 2019). Since its introduction, PHP.B and PHP.eB have been widely used to achieve a broad brain transduction following systemic deliveries. These viral vector capsids are specific for the C57BL/6J mouse strain, and cannot be used in other mouse strains as well as in non-human primates (Hordeaux et al., 2018, 2019; Matsuzaki et al., 2018; Batista et al., 2020; Xie et al., 2021). Compared to PHP.B and PHP.eB, the AAV9-P31 capsid variant exhibited similar performance across different mouse strains tested so far, such as C57BL/6J, Balb/c, and FVB/N (Bunuales et al., 2024; Garcia-Gomara et al., 2025).

### Animal models of Parkinson’s disease based on enhanced neuromelanin accumulation

Although the potential link between NMel and dopaminergic cell vulnerability has long been postulated (Hirsch et al., 1988, 1989; Kastner et al., 1992), this association was often neglected since rodents, as the most commonly used laboratory experimental animals, completely lack NMel pigmentation in SNpc, VTA, and LC neurons. Furthermore, NMel pigmentation of aged dopaminergic cells parallels the accumulation of toxic chemical compounds such as metallic ions that exhibit a high binding affinity with melanins (Karlsson and Lindquist, 2013; Haining and Achat-Mendes, 2017). Furthermore, the chelating nature of NMel allows interaction with neurotoxicants like MPP+ and alpha-synuclein (D’Amato et al., 1986; Wang et al., 2012). In parallel, accumulated clinical evidence supported a bi-directional link between the incidence of Parkinson’s disease and melanoma (reviewed in Flores-Torres et al., 2024). Interestingly, alpha-synuclein expression in malignant melanocytes was recently reported elsewhere (Dean and Lee, 2021).

Existing evidence paved the way for the implementation of novel animal models of PD based on the AAV-mediated enhanced expression of human tyrosinase aimed to induce NMel accumulation in dopaminergic neurons. These approaches were first tested in rats through the intranigral delivery of AAV1-hTyr (Carballo-Carbajal et al., 2019) and later upgraded to non-human primates (Chocarro et al., 2023). Both strategies lead to the characterization of animal models mimicking the known neuropathological hallmarks of PD with unprecedented accuracy, comprising a time-dependent pigmentation of SNpc neurons, presence of intracytoplasmic Lewy body-like aggregates, and a progressive dopaminergic cell death. Moreover, rats injected with AAV1-hTyr displayed an evident motor phenotype (Carballo-Carbajal et al., 2019), whereby a circuit-specific anterograde spread of alpha-synuclein towards the cerebral cortex has been observed in macaques (Chocarro et al., 2023). Although the mechanisms underlying the cross-talk between NMel and alpha-synuclein have not yet been elucidated in detail, these models provided the first demonstration linking NMel accumulation above a given threshold as the key factor leading to the development of an endogenous synucleinopathy in the form of Lewy bodies. In this regard, gene therapy strategies designed to reduce NMel levels have succeeded in preventing the subsequent synucleinopathy and thus leading to dopaminergic cell neuroprotection (Carballo-Carbajal et al., 2019; Gonzalez-Sepulveda et al., 2023). Moreover, the recent introduction of a tissue-specific transgenic mouse model based on the constitutive expression of human tyrosinase represents another step forward in the same direction (Laguna et al., 2024).

### The added value of the PIGMO model

Available animal models of PD have been instrumental in setting up the currently available therapeutic arsenal and obtaining a deeper understanding of the mechanisms underlying the pathophysiological events that typically characterize this neurodegenerative disorder. By definition, each model serves specific research goals, meaning that not a single model fully recapitulates the inherent clinicopathological complexity of PD, and therefore can hardly predict the therapeutic efficacy of novel candidates, such as those with a potential disease-modifying effect (Zeiss et al., 2017; reviewed in Sturchio et al., 2024; see also Barker and Björklund, 2020).

Besides neurotoxin-based animal models, at present the most popular mouse models can be broadly categorized into those related to alpha-synuclein (transgenic, viral-mediated overexpression, preformed fibrils; reviewed in Visanji et al., 2016; Cenci and Björklund, 2020; Zhang et al., 2022; Björklund and Mattsson, 2024) and those focused on highly penetrant rare genetic mutations linked to familial forms of PD, such as those related to the PARK family, LRRK2, PINK1, GBA1, and DJ-1 (reviewed in Bastioli et al., 2021; Shadrina and Slominsky, 2021; Lal et al., 2024). Mouse models outside these categories are the Mitopark mouse (Ekstrand et al. 2007), as well as those being recently introduced, such as the one based on loss of functional mitochondrial complex I (Gonzalez-Rodriguez et al., 2021) and the transgenic mouse line M12 (Wegrzynowicz et al., 2019). In general terms, and regardless of the chosen rodent model of PD, not a single model managed to mimic the real human disorder, however, these models have a proven record of success when investigating specific disease mechanisms. For instance, viral enhancement of alpha-synuclein expression in the SNpc often resulted in a non-consistent dopaminergic degeneration (Garcia Moreno et al., 2024).

Accordingly, for most of the mouse models listed above, the PIGMO model introduced here can be regarded as complementary instead of an alternative model. In other words, the intravenous delivery of AAV9-P31-hTyr can easily be used as a feasible strategy enabling a second insult intended to boost the PD phenotype in models lacking the right outcomes. Moreover, the PIGMO model has several advantages, such as (i) simplicity of implementation by merely needing a systemic administration of the viral vector encoding the human tyrosinase gene leading to a bilateral PD model without the need to perform stereotaxic surgeries for intraparenchymal injections, (ii) a wide therapeutic window enabling testing disease-modifying therapeutic interventions, (iii) a highly predictable and reproducible pattern of neuropathological and motor readouts, enabling administration of therapeutics once the neurodegenerative processes are already ongoing but before reaching a non-returning point.

## Supporting information

Plasmid map and sequence for pAAV-CMV-hTyr and pAAV-CMV-hTyr:STOP

Tier-specific pigmentation of dopaminergic neurons

Lewy body-like inclusions in LC neurons

Cellular phenotypes SNpc & VTA

Pigmented peripheral organs

## FUNDING

This research was funded in whole or in part by Aligning Science Across Parkinson’s (Grant No. ASAP-020505) through the Michael J. Fox Foundation for Parkinson’s Research (MJFF). For the purpose of open access, the author has applied a CC-BY 4.0 public copyright license to all Author Accepted Manuscripts arising from this submission. Work was also funded by MICIU/AIE/10.13039/501100011033 (Grant No. PID2023-127802OB-I00) and FEDER, UE.

## COMPETING INTERESTS

The authors report no competing interests.

## DATA AVAILABILITY

Further information and requests for resources and reagents should be directed to and will be fulfilled by the corresponding author, Jose L. Lanciego. The data, code, protocols, and key lab materials used and generated in this study are listed in a Key Resource Table alongside their persistent identifiers at https://doi.org/10.5281/zenodo.15386050. No code was generated for this study; all data cleaning, preprocessing, analysis, and visualization were performed using Fiji ImageJ, GraphPad Prism 9.0.2, Stata 14, and Aiforia.

## Notes

https://doi.org/10.5281/zenodo.15386050

### Competing Interest Statement

The authors have declared no competing interest.

### Summary of Updates

KRT table updated, reference to the new version of the KRT added to the manuscript

