## Supplementary figures and images for "Introducing PIGMO, a novel PIGmented MOuse model of Parkinson’s disease (V1)"

### Cellular phenotypes SNpc & VTA

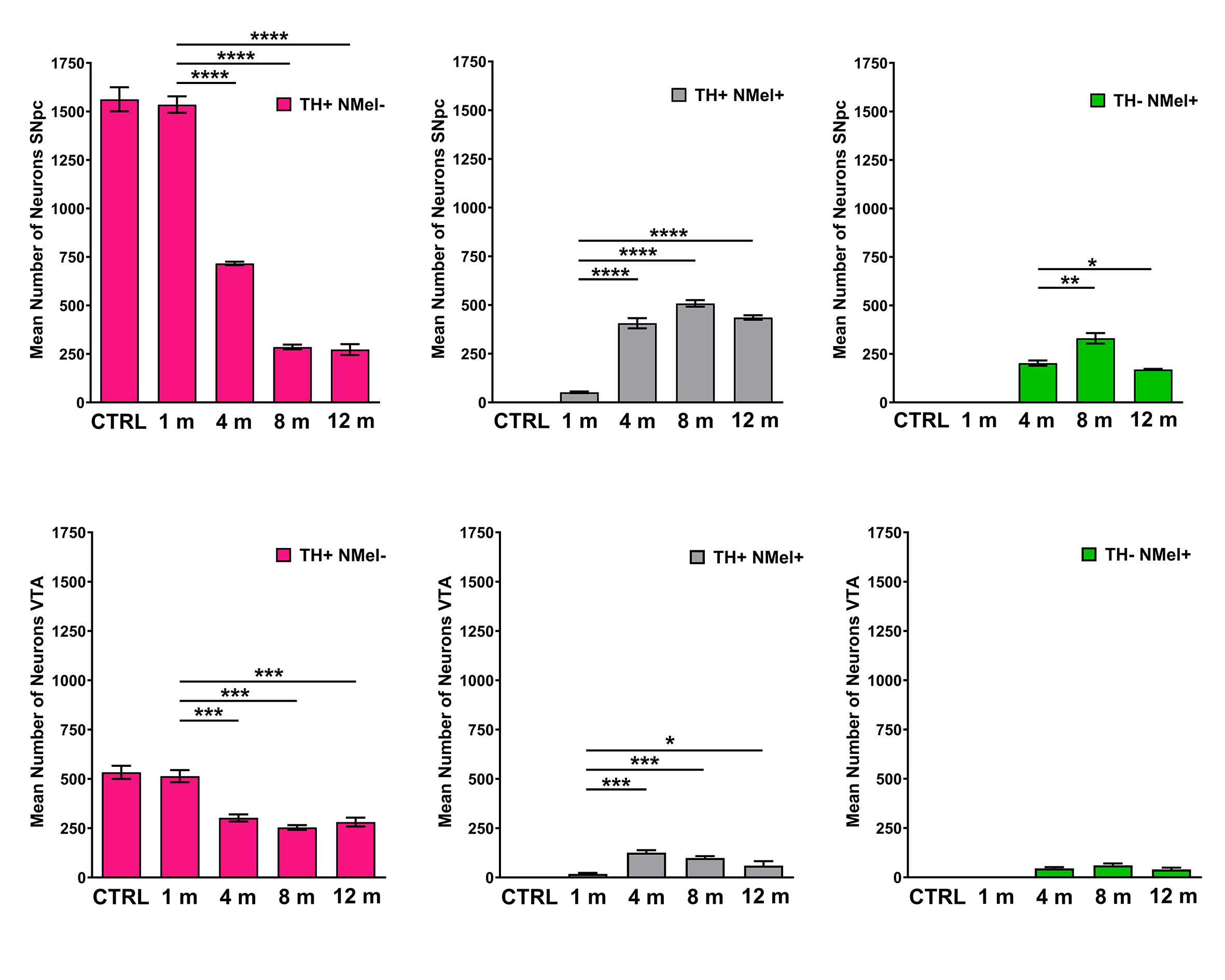

### Lewy body-like inclusions in LC neurons

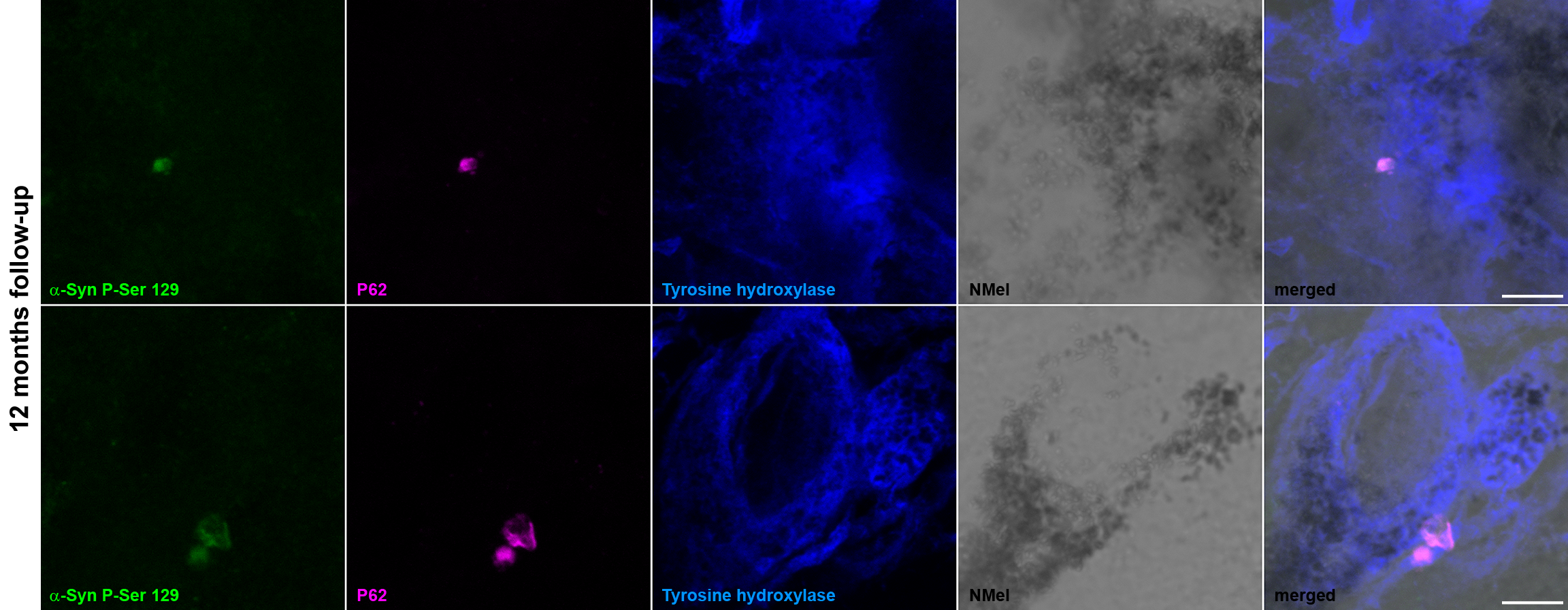

### Pigmented peripheral organs

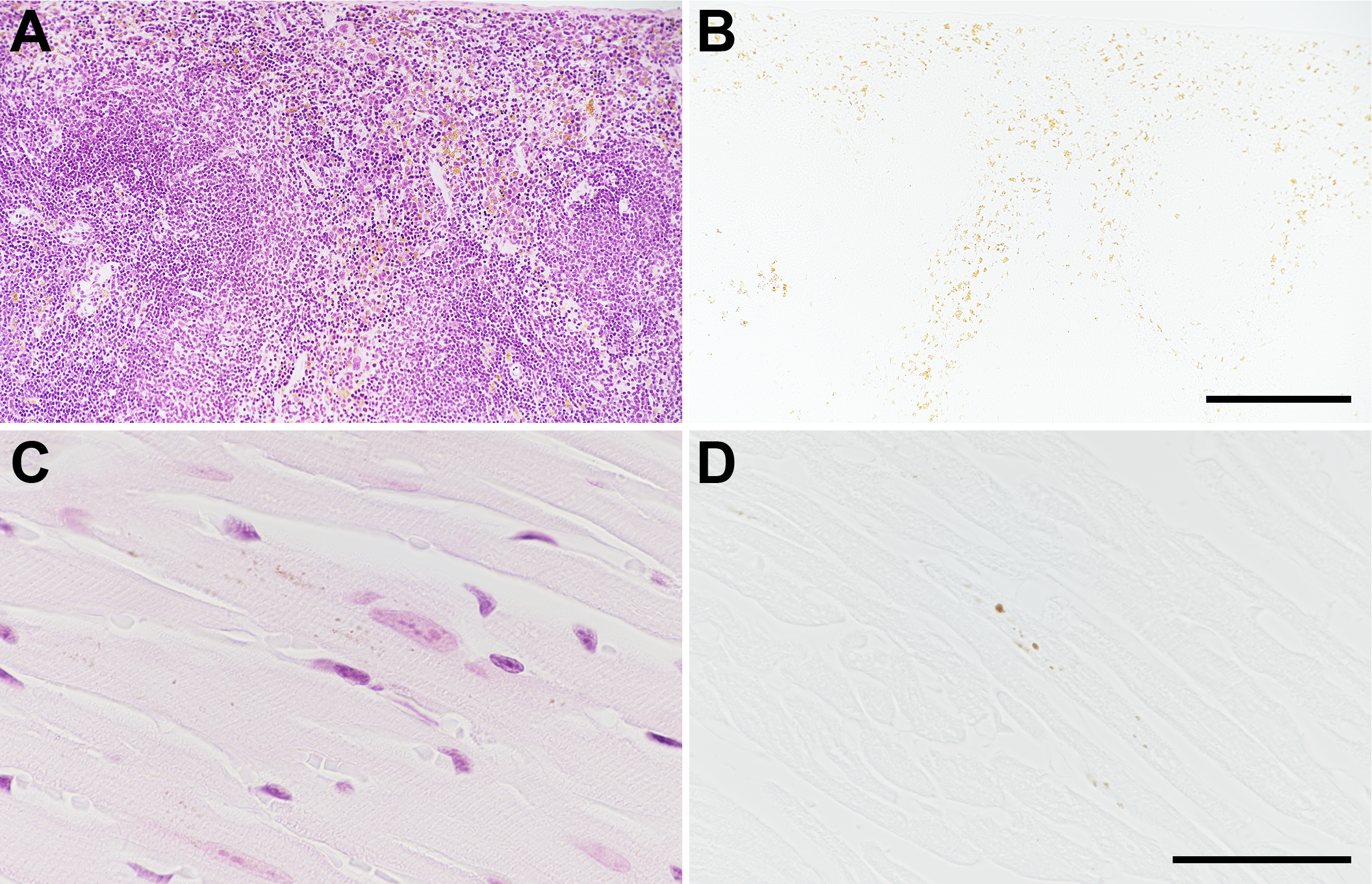

### Plasmid map and sequence for pAAV-CMV-hTyr and pAAV-CMV-hTyr:STOP

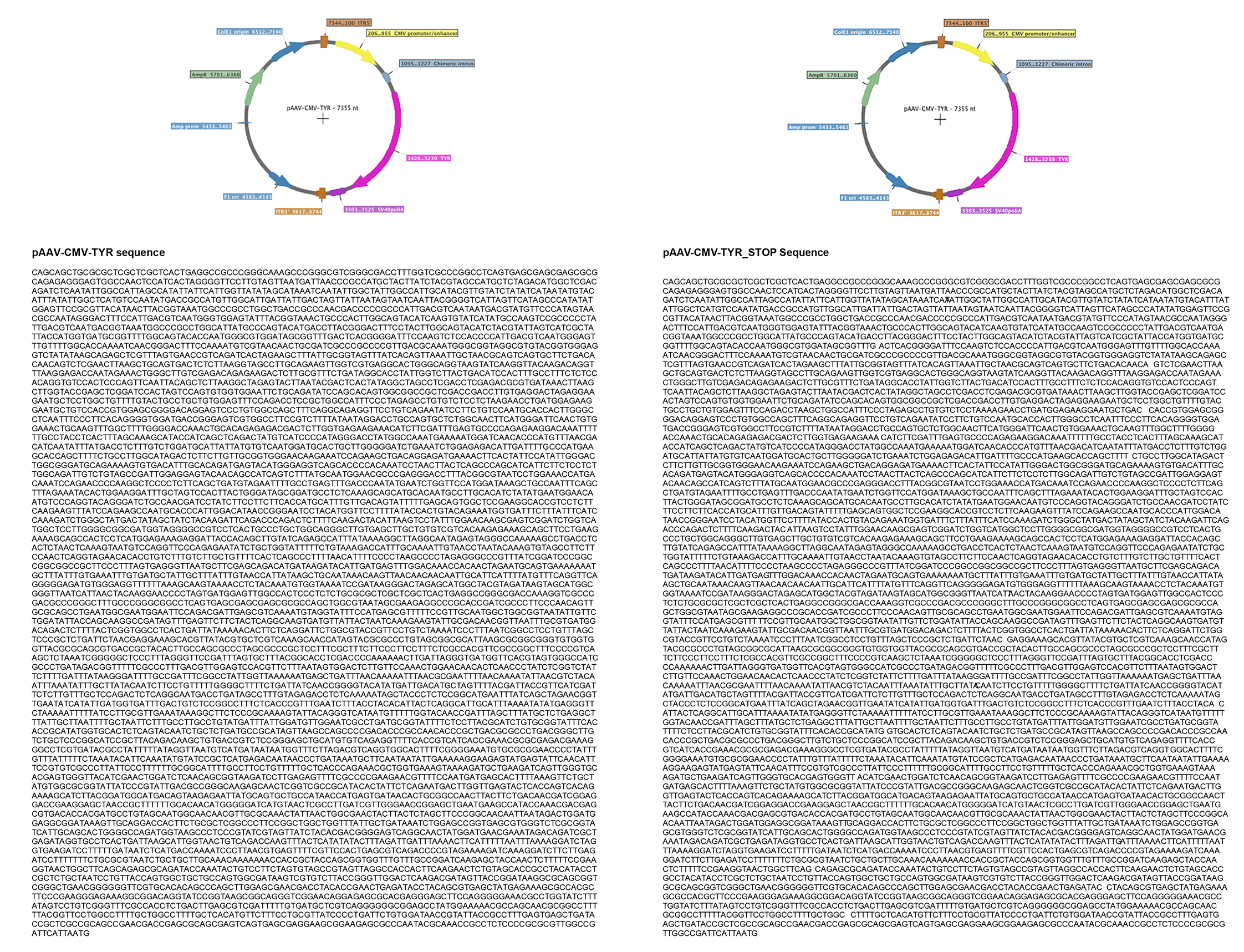

### Tier-specific pigmentation of dopaminergic neurons

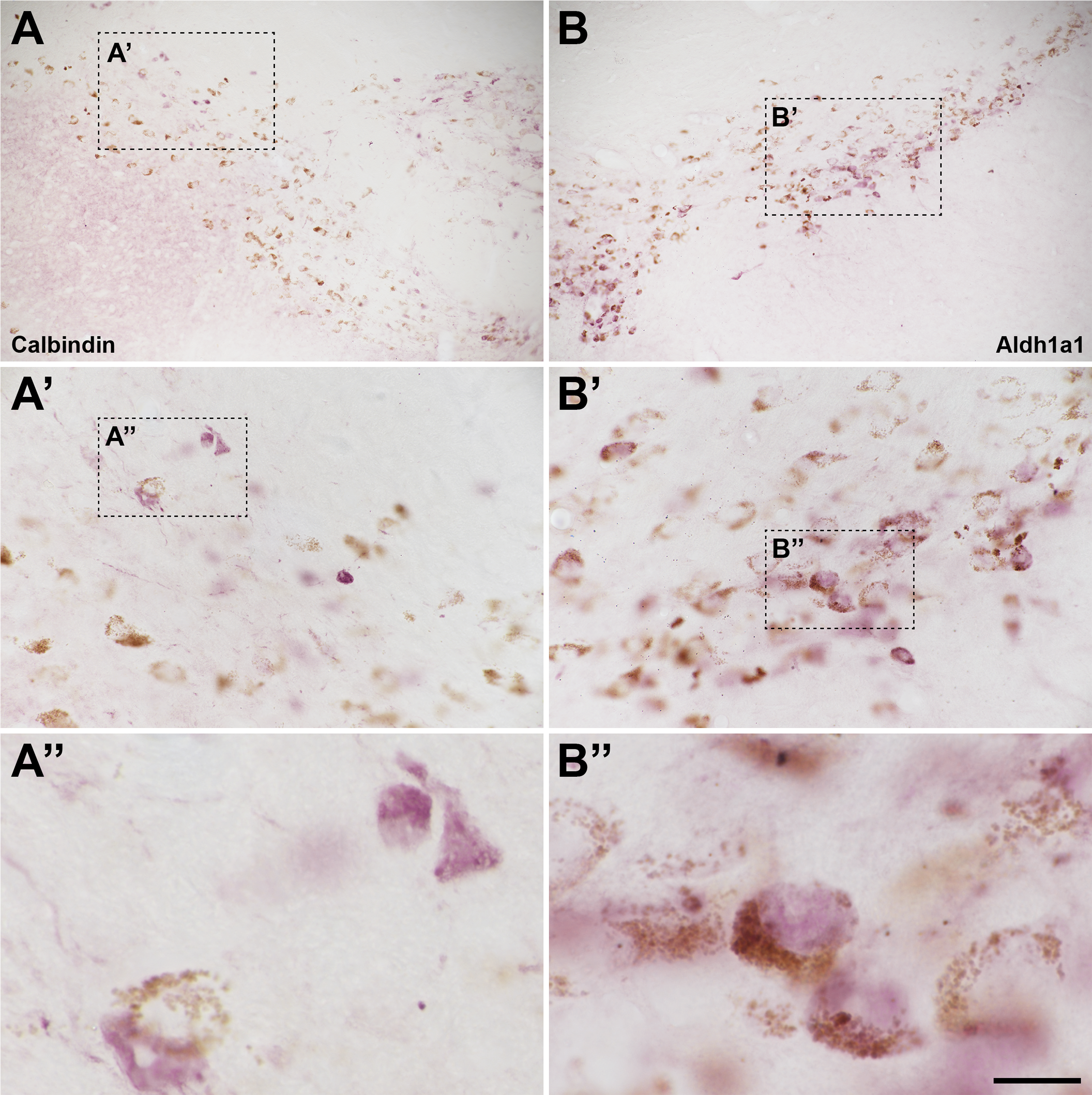
